# Pacsin 2-dependent N-cadherin internalization regulates the migration behaviour of malignant cancer cells

**DOI:** 10.1101/2022.08.08.502718

**Authors:** Haymar Wint, Jianzhen Li, Tadashi Abe, Hiroshi Yamada, Yasutomo Nasu, Masami Watanabe, Kohji Takei, Tetsuya Takeda

## Abstract

Cell migration is essential for both physiological and pathological processes such as embryonic morphogenesis, wound repair and metastasis of cancer cells. Collective cell migration is the coordinated movement of multiple cells connected with cadherin-based adherence junctions. Cadherins undergo dynamic intracellular trafficking and their surface level is determined by a balance between endocytosis, recycling and degradation. However, regulatory mechanisms of cadherin turnover in the collective cell migration remain to be elucidated.

In this study, we show that a BAR domain protein pacsin 2 plays an essential role in collective cell migration by regulating the internalization of N-cadherin in human bladder cancer cells T24. Pacsin 2 and its associating GTPase dynamin 2 colocalized with N-cadherin at the cell periphery in T24 cells. Depletion of either pacsin 2 or dynamin 2 induced interdigitating cell-cell contacts enriched with N-cadherin. Imaging analyses of the wound healing assay showed that pacsin 2-depleted T24 cells migrated in a collective and directed manner in contrast with randomly migrating control cells. Furthermore, cell-surface biotinylation assay showed that internalization of N-cadherin is attenuated in pacsin 2-depleted cells. Interestingly, the GST-pulldown assay demonstrated that the SH3 domain of pacsin 2 binds to the cytoplasmic domain of N-cadherin, suggesting that surface levels of N-cadherin are regulated by pacsin 2-mediated endocytosis. These data support new insights into a novel endocytic route of N-cadherin in collective cell migration providing pacsin 2 as a possible therapeutic target for cancer metastasis.

## INTRODUCTION

Cell migration is fundamental for diverse physiological and pathological processes including development, immune response and cancer metastasis (Yamada and Sixt, 2019). Cancer cells migrate either individually or collectively during metastasis (Pandya et al., 2017). Collectively migrating cancer cells are generally more aggressive and resistant to chemotherapies compared to individually migrating cancer cells (Aceto et al., 2014). Collective cell migration is a coordinated movement of a group of cells that are connected via adherence junctions (Friedl and Gilmour, 2009; Rorth, 2009). Different guidance mechanisms such as chemotaxis, haptotaxis, durotaxis, and strain-induced mechanosensing are involved in the collective movement of cells (Haeger et al., 2015; Shellard and Mayor, 2021). For successful collective cell migration, two groups of cell adhesion molecules play essential roles in generating and coordinating mechanical forces among cells: focal adhesion molecules such as integrins that transmit forces between cells and underlying ECM, and adherence junction molecules such as cadherins that transmit forces at intercellular adhesion sites (Halbleib and Nelson, 2006; Ray et al., 2017).

Cadherins are homophilic calcium-dependent cell adhesion molecules that play important roles in various physiological and pathological processes such as development (Gumbiner, 2005; Halbleib and Nelson, 2006) and cancer (Kaszak et al., 2020). There are over 100 different cadherin subtypes in vertebrates that are classified into four groups: classical, desmosomal, protocadherins and unconventional cadherins (Yagi and Takeichi, 2000). Each cadherin contains a large extracellular ectodomain followed by a transmembrane domain and a small cytoplasmic domain from N- to C-terminus (Oda and Takeichi, 2011). Interactions of ectodomains from apposed cells mediate cell-cell contact, whereas the cytoplasmic domain contributes to linking cadherins to the underlying actin cytoskeleton by forming a complex with α- and β-catenins (Ratheesh and Yap, 2012). The cytoplasmic domain of cadherins also binds to p120-catenin (p120) which controls cadherin endocytosis and turnover thus regulating cell surface cadherin levels responsible for cell-cell adhesion (Cadwell et al., 2016). A recent study showed that classical cadherins, E- and N-cadherins, mediate cell-cell contacts to enhance the spreading efficiency of the collectively migrating cells (Zisis et al., 2021). Another study on collectively migrating endothelial cells showed that polarized membrane protrusions enriched with unconventional VE-cadherin called “cadherin fingers” serve as guidance cues that direct collective cell migration (Hayer et al., 2016). Furthermore, classical P-cadherin enhances collective cell migration of myoblasts by activating Cdc42 increasing the strength and anisotropy of mechanical forces (Plutoni et al., 2016). The cadherin-mediated cell-cell contact is determined by a balance between endocytosis, recycling, and degradation (Akhtar and Hotchin, 2001; Cadwell et al., 2016; Kowalczyk and Nanes, 2012; Le et al., 1999). However, the regulatory mechanisms of cadherin turnover in collective cell migration remain to be elucidated.

Bin/amphiphysin/Rvs (BAR) domain proteins are conserved protein families that possess deformation and sensing of membrane curvature (Peter et al., 2004; Safari and Suetsugu, 2012). BAR domain proteins play crucial roles in endocytosis (Takei et al., 1999), exocytosis (Pinheiro et al., 2014), cell migration (Sanchez-Barrena et al., 2012), cytokinesis (Takeda et al., 2013), and cancer metastasis (Yamamoto et al., 2011). BAR domains form “crescent-shaped” dimers that are classified into three subtypes with distinctive topology and curvature: N-BAR (N-terminal amphipathic helix and BAR), F-BAR (Fes/CIP4 homology-BAR) and I-BAR (Inverse-BAR) (Qualmann et al., 2011; Safari and Suetsugu, 2012). Pacsin (protein kinase C and casein kinase substrate in neuron) or syndapin (synaptic dynamin-associated protein) contains an F-BAR domain and an SH3 domain in its N- and C-termini, respectively (Dumont and Lehtonen, 2022). Three pacsin isoforms are expressed in mammalian cells: the neuronal isoform pacsin 1, the muscle-specific isoform pacsin 3, and the ubiquitously expressed isoform pacsin 2 (Modregger et al., 2000). Pacsin 2 has been implicated in caveolar endocytosis, vesicle trafficking, actin dynamics, and cell migration (Chandrasekaran et al., 2016; de Kreuk et al., 2011; Hansen et al., 2011; Senju et al., 2011). Previous studies showed that dynamin 2, a major pacsin 2-associated protein, is required for the internalization of E-cadherin (Miyashita and Ozawa, 2007; Paterson et al., 2003) and VE-cadherins (Chiasson et al., 2009). However, the requirement of pacsins in cadherin turnover remains elusive.

In the present study, we show that pacsin 2 is involved in the collective cell migration of cancer cells by controlling N-cadherin internalization. Depletion of pacsin 2 in T24 bladder cancer cells induces clustering of cells with N-cadherin enriched cell-cell contacts. Electron microscopy shows that the cell-cell contacts induced by pacsin 2 depletion consist of interdigitating finger-like membranous protrusions. Imaging analyses of wound healing assay demonstrate that pacsin 2-depleted cells exhibit a cell migration in a directed manner. Furthermore, cell surface biotinylation and endocytosis assay shows that N-cadherin internalization is inhibited in pacsin 2-depleted T24 cells. Interestingly, the GST pulldown assay shows that the SH3 domain of pacsin 2 binds to the cytoplasmic domain of N-cadherin, suggesting the direct roles of pacsin 2 in regulating N-cadherin endocytosis that affects cell migration behaviour of malignant cancer cells.

## RESULTS

### Pacsin 2 localizes at the cell periphery in T24 cells

To determine the functions of pacsins in malignant cancer cells, expression and localization profiles of pacsin isoforms were examined in T24 cells. Immunoblot analysis of the whole-cell extract from T24 cells exhibited that pacsin 1 and pacsin 2, but not pacsin 3, were expressed in T24 cells (Fig.S1, A). Immunofluorescence microscopy showed that pacsin 1 dispersedly localized in the cytoplasm, whereas pacsin 2 localized to the cell periphery in T24 cells (Fig.S1, B). Pacsin 2 interacts with dynamin 2 which is required for invadopodia formation of T24 cells (Zhang et al., 2016). However, pacsin 2 did not colocalize with dynamin 2 at the perinuclear invadopodia, but they colocalized at the cell periphery (Fig. 1). Localization of pacsin 2 to the cell periphery was also confirmed by observing that cortactin, an essential actin organizer at the invadopodia, was not colocalized with pacsin 2 (Fig. 1). These results suggest that, unlike dynamin 2, pacsin 2 is not involved in invadopodia formation but it plays a role at the cell periphery such as cell migration.

**Fig. 1.**
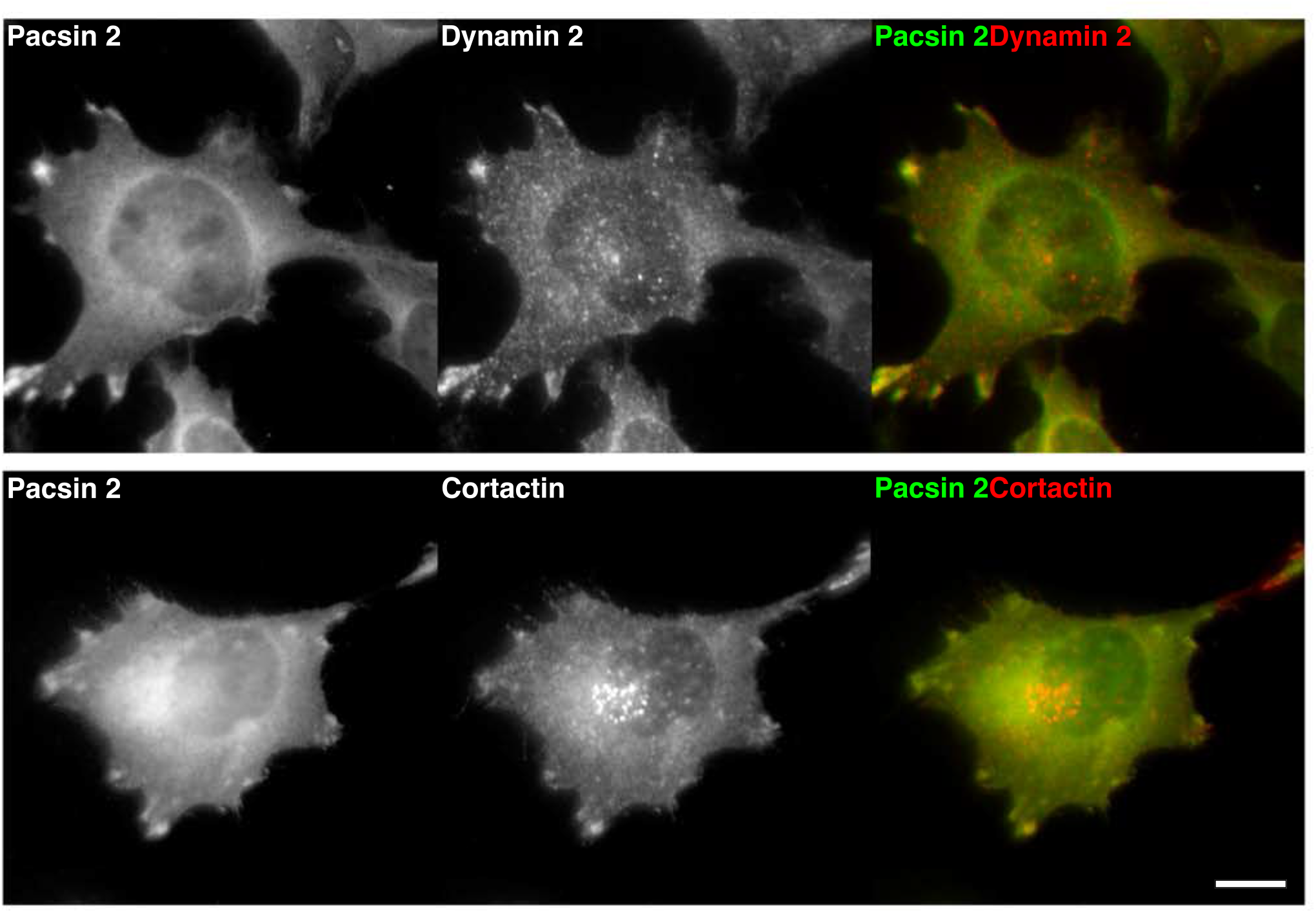
Pacsin 2 localises to the cell periphery in T24 cells. Immunofluorescence microscopic images of endogenous pacsin 2 (green) with either endogenous dynamin 2 (red) or endogenous cortactin (red), and their merged images. The scale bar is 10 μm.

### Pacsin 2 depletion enhances the migration of T24 cells

To elucidate if pacsin 2 is involved in the migration of T24 cells, the effect of pacsin 2- depletion was examined in the wound healing assay. Control RNAi cells of T24 migrated slowly and only 15.7% of the scratched area was filled in 12 hours (Fig. 2, A and B, siCtrl). In contrast, pacsin 2 RNAi cells showed enhanced migration activity and the wound closure area was extended to 31.3 - 54.1% in 12 hours (Fig. 2, A and B, siPacsin 2 #1-3). Immunoblot analyses confirmed that all three different siRNAs targeting pacsin 2 caused efficient depletion of pacsin 2 (Fig. 2, C). These results suggest that pacsin 2 negatively regulates cell migration activities of T24 cells.

**Fig. 2.**
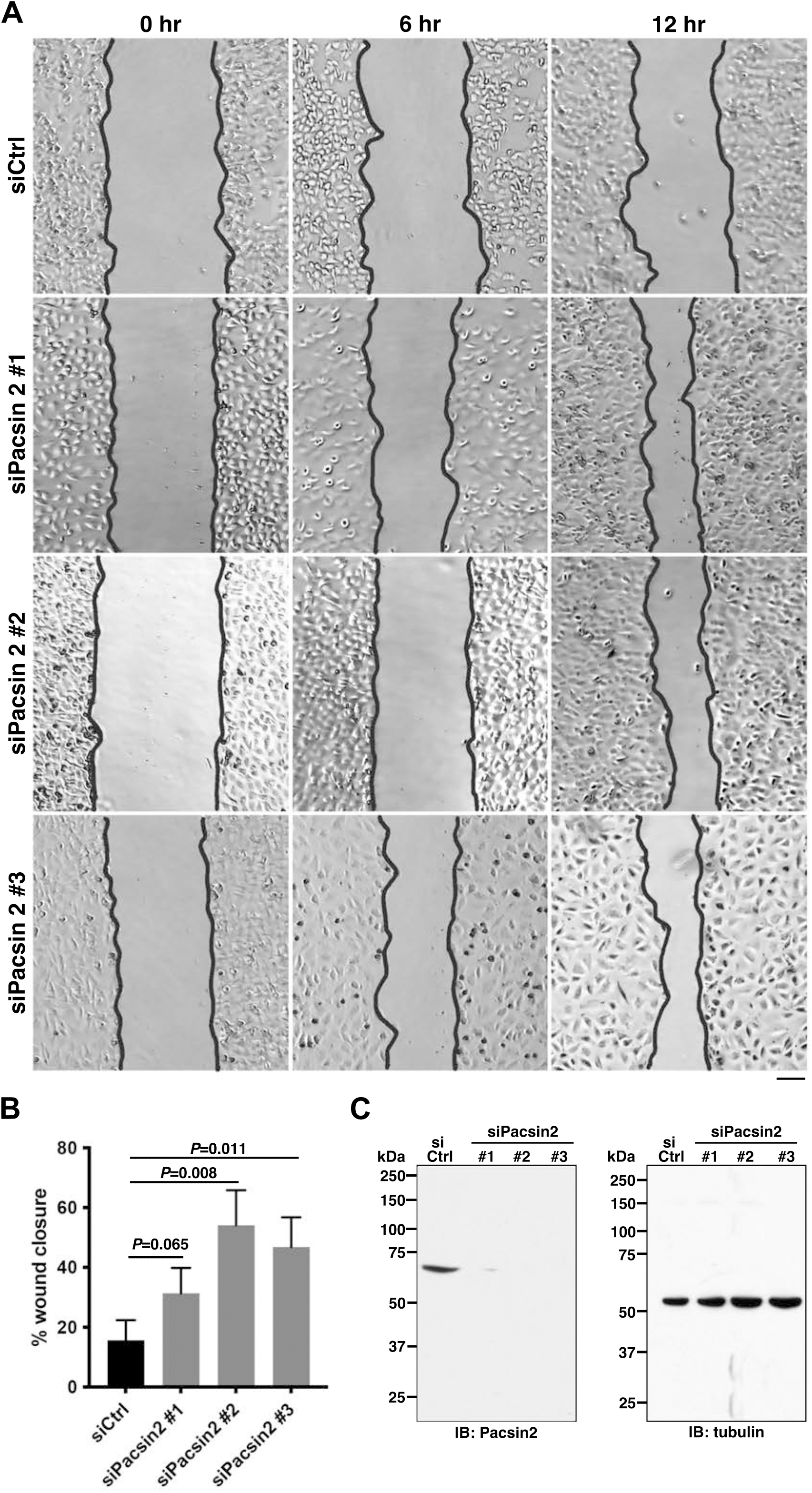
Pacsin 2 depletion enhances cell migration in T24 cells. (A) DIC images of migrating cells in the wound healing assay. Representative micrographs show either control RNAi cells (siCtrl) or pacsin 2 RNAi cells (siPacsin 2 #1, #2 and #3) at 0, 6 and 12 hours after the start of the wound healing assay. The scale bar is 200 μm. (B) Quantitation of wound closure by either control RNAi cells (siCtrl) or pacsin 2 RNAi cells (siPacsin 2 #1, #2 and #3). Data are means ± SD. (C) Immunoblot analysis of cell extract from control RNAi cells (siCtrl) or pacsin 2 RNAi cells (siPacsin 2 #1, #2 and #3) using antibodies against pacsin 2 (IB: Pacsin 2) or tubulin as an internal control (IB: tubulin).

To clarify the cause of the enhanced cell migration by pacsin 2 RNAi cells, the dynamics of cell migration were analyzed by live-cell imaging of the wound healing assay. Tracking of representative cells showed that control RNAi cells moved with an average speed of 4.1 *μ*m/min, but they moved randomly (Fig.3, A-D, siCtrl; Movie S1). In contrast, pacsin 2 RNAi cells migrated in a more directed manner, though their velocity was comparable to that of control RNAi cells (3.5 *μ*m/min), (Fig. 3, A-D, siPacsin 2; Movie S2). These results suggest that pacsin 2 has a role in regulating the direction of cell migration.

**Fig. 3.**
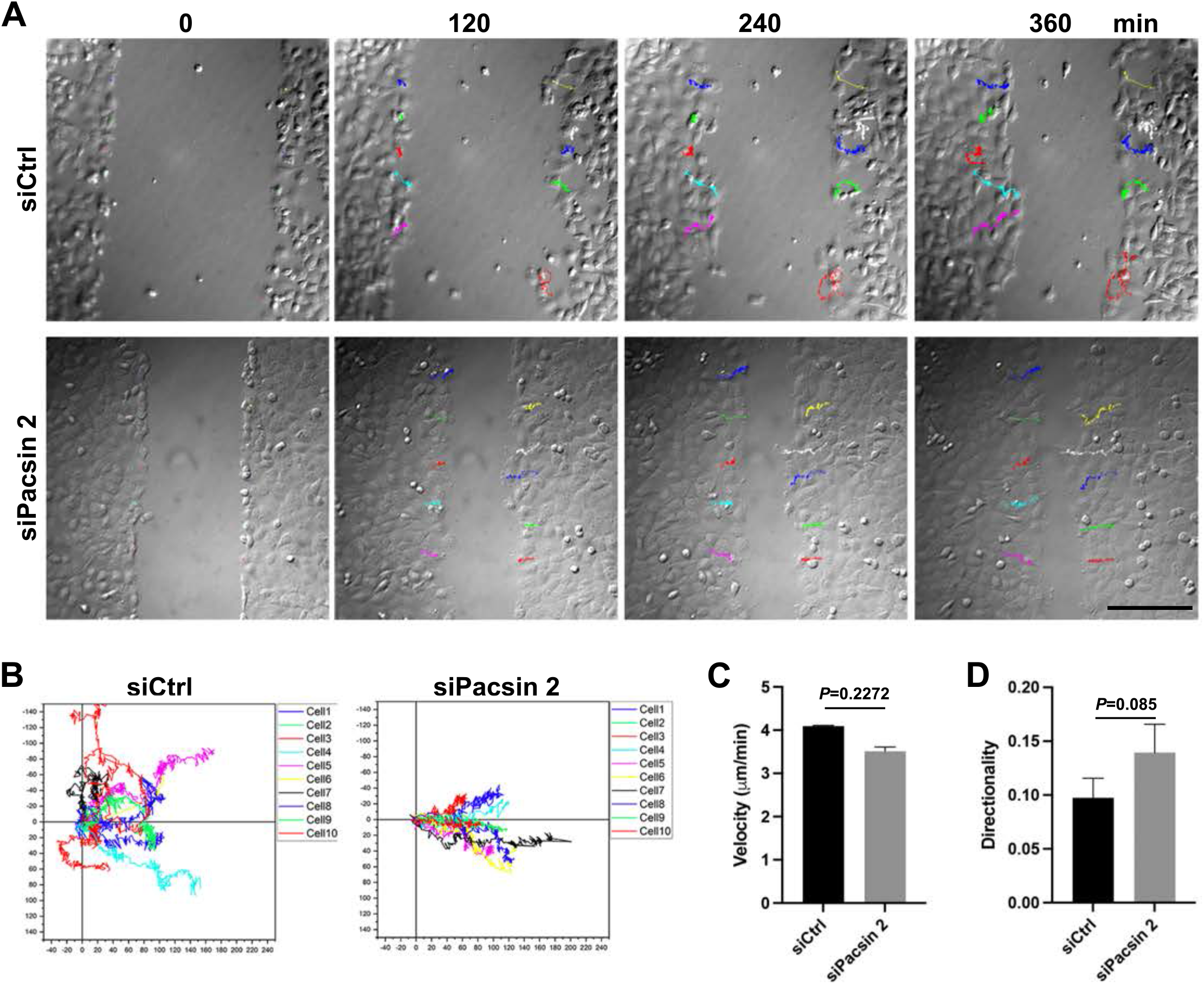
Pacsin 2 depletion induces directional cell migration. (A) Live-cell imaging analysis of the wound healing assay. Time-lapse images of control RNAi (siCtrl) or pacsin 2 RNAi (siPacsin 2) cells at 0, 120, 240 and 360 min after the start of wound healing assay. Traced paths of ten representative cells were shown in different colours. (B) Trajectories of cell tracking for representative cells over 360 min. (C) Quantitation of cell velocity in the wound healing assay. Data are means ± SD (n=10 cells, N=3) in 360 min of the wounding healing assay. (D) Quantitative analysis of cell directionality in the wound healing assay. Data are means ± SD (n=10 cells, N=3) in 360 min of the wounding healing assay.

### Pacsin 2 depletion induces cell-cell contacts enriched with N-cadherin

To elucidate how pacsin 2 can affect the directionality of cell migration, cellular phenotypes were analyzed by immunofluorescence microscopy. Control RNAi cells tended to grow individually and only 34.5% of cells form cell-cell contacts at subconfluent cell densities (Fig. 4, A and B, siCtrl). In contrast, pacsin 2 RNAi induced cell clustering and more than 77.5% of cells exhibited cell-cell contacts (Fig. 4, A and B, siPacsin 2 #1, 2 and 3). Similarly, cell cluster formation was also induced by dynamin 2 RNAi (55.3%), while only 26.4% of control RNAi cells exhibited cell-cell contacts (Fig. S2, A and B). These results suggest that pacsin 2 and dynamin 2 are involved in the formation of cell-cell contacts in T24 cells.

**Fig. 4.**
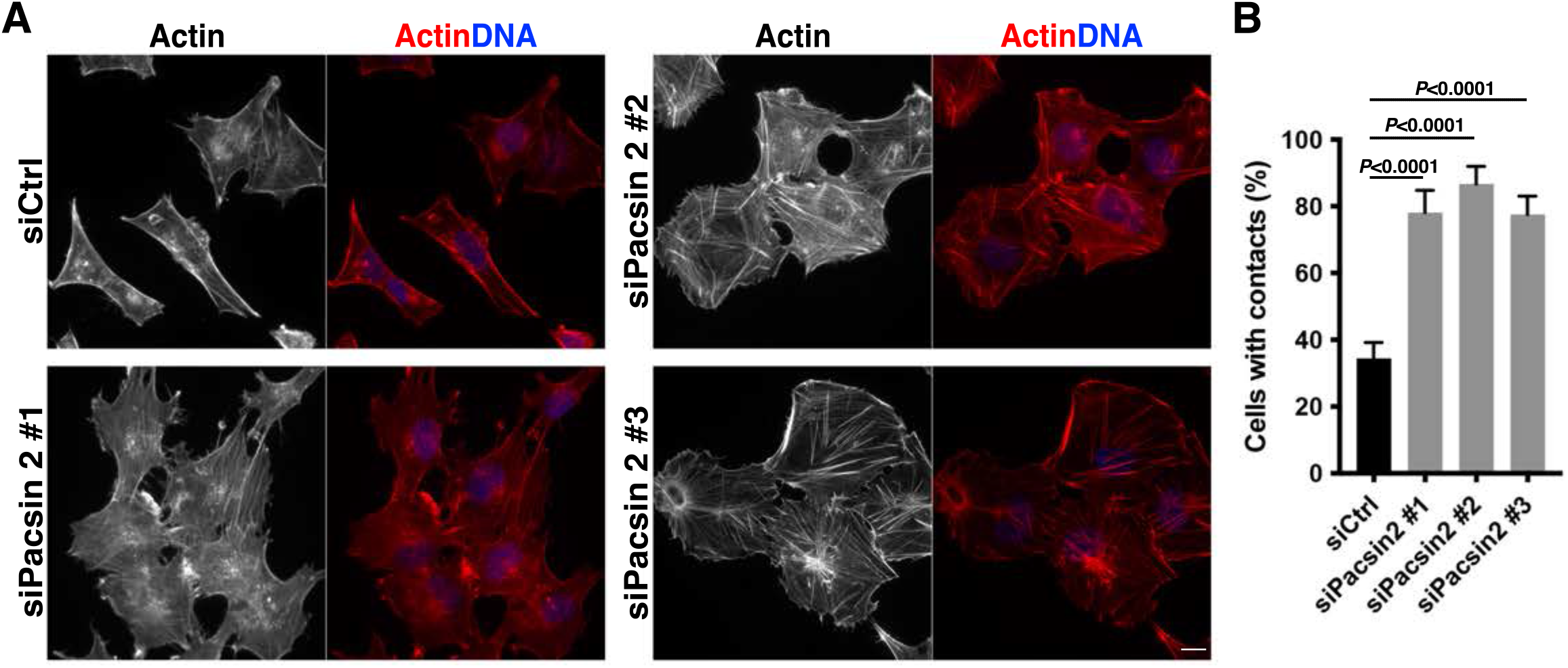
Depletion of pacsin 2 induces cell-cell contacts in T24 cells. (A) Immunofluorescence micrographs of control RNAi cells (siCtrl) or pacsin 2 RNAi cells (siPacsin 2 #1, #2 and #3) stained for F-actin (red) and its merged images with DNA (blue). The scale bar is 10 μm. (B) Quantitation of cells with cell contacts in control RNAi cells (siCtrl) or pacsin 2 RNAi cells (siPacsin 2 #1, #2 and #3). Data are means ± SD (n≥120 cells, N=3).

To identify the molecular components of the cell-cell contacts induced in pacsin 2 or dynamin 2 RNAi cells, the expression and localization profiles of cadherins were examined. Immunoblot analyses showed that RT4, which represents papillary bladder carcinoma, expressed E-cadherin but not N-cadherin, whereas T24 which represents more aggressive bladder cancer mainly expressed N-cadherin but not E-cadherin (Fig. S3). In contrast, unconventional VE-cadherin or classical P-cadherin were not expressed in T24 cells (Fig. S3). To determine if N-cadherin is a component in the cell-cell contacts induced in pacsin 2 or dynamin 2 RNAi cells, the localization of N-cadherin in T24 cells was analyzed by immunofluorescence microscopy. Endogenous N-cadherin colocalized with either pacsin 2 or dynamin 2 at the cell periphery in T24 cells (Fig. S4). Similarly, N-cadherin localized at the cell periphery as well as cytoplasmic dots in control RNAi cells (Fig. 5, siCtrl). In contrast, in pacsin 2 RNAi cells, N-cadherin was accumulated at the cell-cell contact sites where actin filaments extended from contacting cells are interdigitating (Fig. 5, siPacsin2 #1-3). A similar distribution of N-cadherin to the cell-cell contact sites was also observed in dynamin 2 RNAi cells (Fig. S5) suggesting their functional association in the formation of N-cadherin-rich cell-cell contacts.

**Fig. 5.**
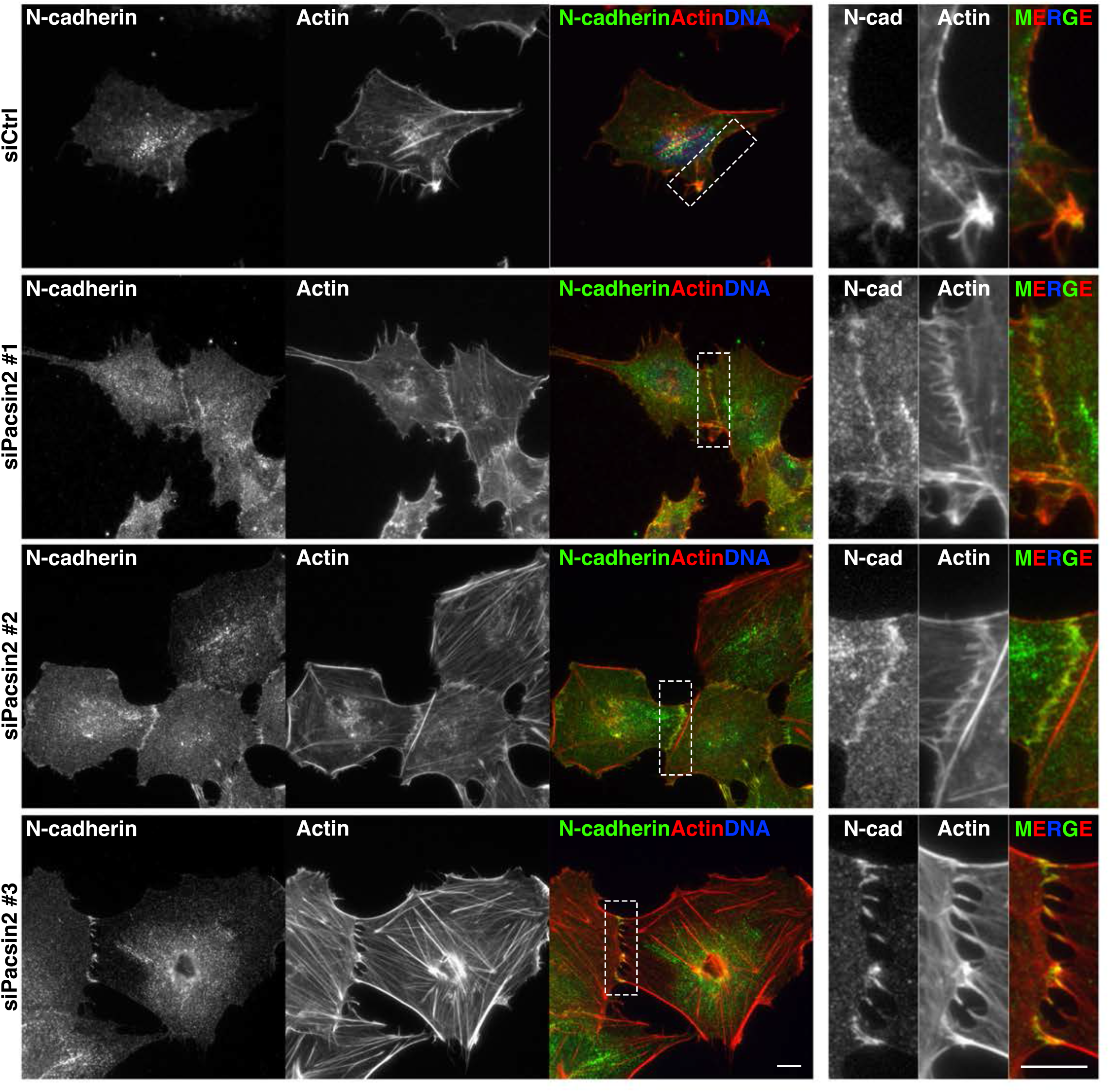
Pacsin 2 depletion induces N-cadherin-rich cell-cell contacts. Immunofluorescence micrographs of control RNAi cells (siCtrl) and pacsin 2 RNAi cells (siPacsin 2 #1, #2 and #3) stained for endogenous N-cadherin (green), F-actin (red) and their merged images with DNA (blue). Enlarged images show either the cell periphery in control cells or N-cadherin-rich cell-cell contact sites in pacsin 2 RNAi cells (shown in the dashed rectangle). Scale bars are 10 μm.

To gain further insights into the cell-cell contact sites induced by pacsin 2 RNAi, their ultrastructure was analyzed by electron microscopy. Control RNAi cells sometimes form cell-cell contacts, but the plasma membrane structures between closely apposed cells were smooth (Fig. 6, siCtrl). In contrast, in pacsin 2 RNAi cells, numerous membranous protrusions were formed at the cell-cell contact sites and they were often interdigitating each other (Fig. 6, siPacsin 2). Importantly, immunoblot analyses showed that the total amount of N-cadherin was not altered in either pacsin 2 RNAi cells (Fig. S6, A) or dynamin 2 RNAi cells (Fig. S6, B), suggesting that the cell surface level of N-cadherin, but not its transcription and/or translation, was regulated by pacsin 2 or dynamin 2 in T24 cells to induce cell-cell contacts.

**Fig. 6.**
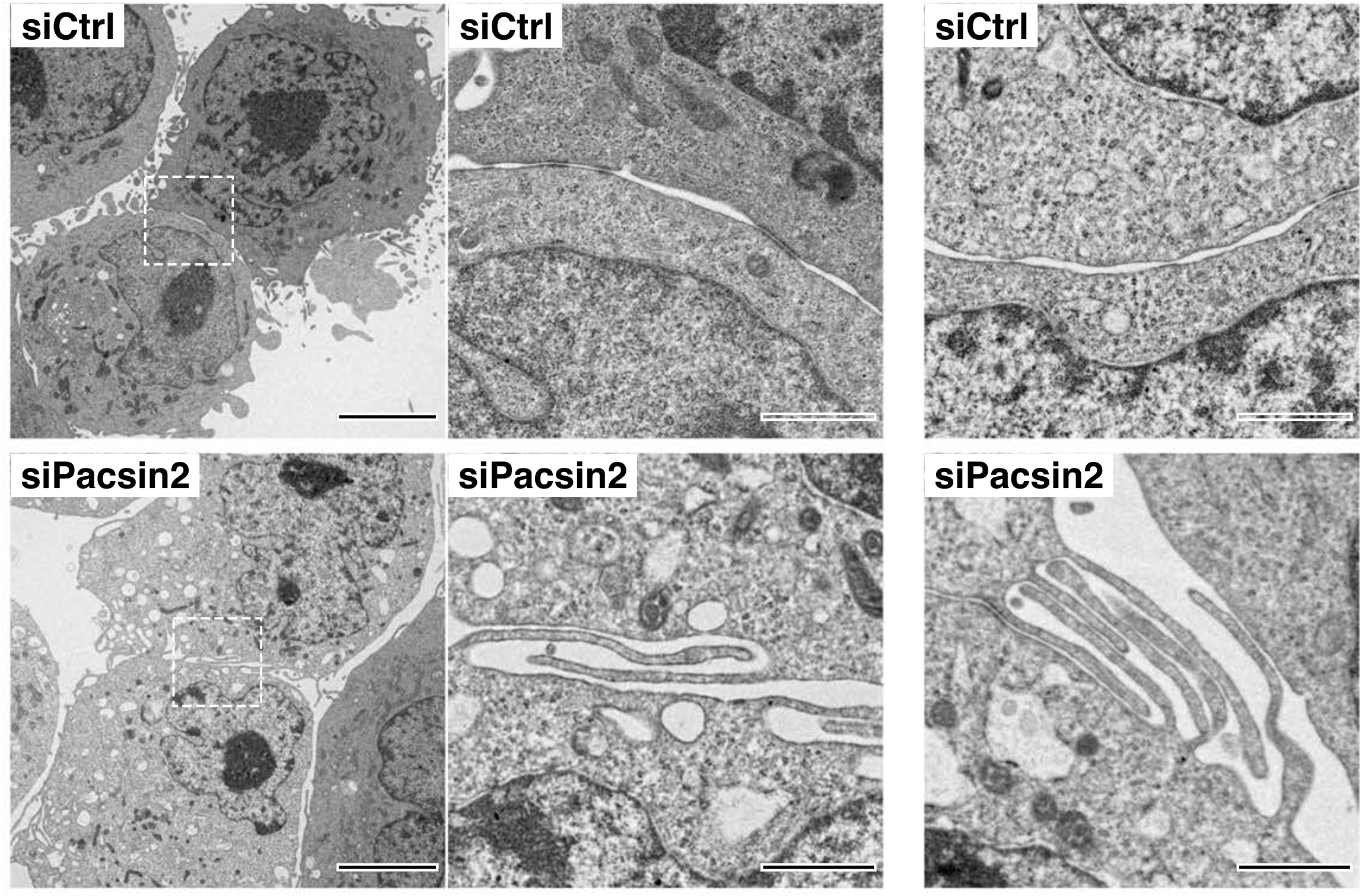
Pacsin2 depleted cells form interdigitating membrane protrusions at the cell-cell contact sites. Transmission electron microscopic images of cell-cell contact sites in control RNAi cells (siCtrl) and pacsin 2 RNAi cells (siPacsin 2) at different magnifications (0.7K and 4K). Scale bars are 5 μm for 0.7K and 1 μm for 4K.

### Depletion of pacsin 2 attenuates N-cadherin endocytosis in T24 cells

The cell surface level of cadherin is determined by a balance between endocytosis, recycling and degradation. Since both pacsin 2 and dynamin 2 have been implicated in endocytosis, the internalization of surface N-cadherin was analyzed by surface biotinylation and endocytosis assay. In both control and pacsin 2 RNAi cells, N-cadherin on the cell surface was internalized within 30 min and overall internalized N-cadherin level was gradually decreased probably due to degradation (Fig. 7, A and B). However, in pacsin 2 RNAi cells, internalization of N-cadherin was attenuated and the relative amount of internalized N-cadherin was around 46.3% of the control cells at 30 min after the restart of endocytosis (Fig. 7, A and B). To address if pacsin 2 plays a direct role in regulating N-cadherin endocytosis, the interaction between pacsin 2 and N-cadherin was examined using GST pull-down assay. Pacsin 2 contains a C-terminal SH3 domain that binds to proline-rich motifs in its interacting proteins (Fig. 8, A). Interestingly, the cytoplasmic domain of N-cadherin contains two PxxP motifs that potentially bind to the SH3 domain of pacsin 2 (Fig. 8, B). Indeed, the GFP-tagged cytoplasmic domain of N-cadherin is bound to the GST-tagged pacsin 2 SH3 domain, but not with GST bait (Fig. 8, C), suggesting that pacsin 2 has a direct role in regulating N-cadherin internalization.

**Fig. 7.**
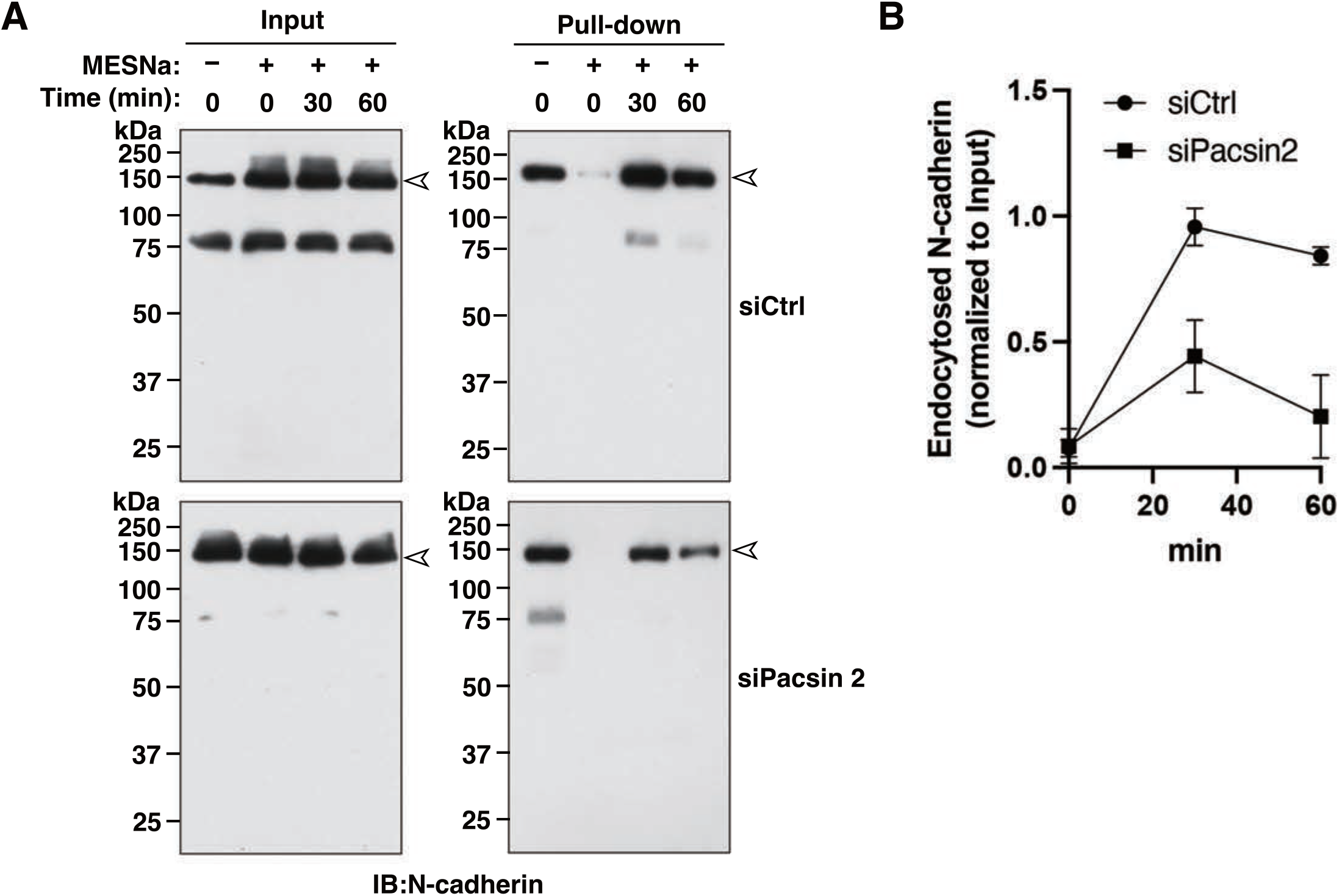
Pacsin 2 is required for the internalization of N-cadherin. (A) Representative immunoblots of N-cadherin internalization experiments. Surface N-cadherin in control RNAi cells (siCtrl) and pacsin 2 RNAi cells (siPacsin 2) were biotinylated at 4℃ and incubated at 37 °C for the indicated periods to allow endocytosis. Immunoblots for total N-cadherin (Input) and internalized N-cadherin (Pull-down) in control RNAi cells (siCtrl) or pacsin 2 RNAi cells (siPacsin 2) are shown. (B) Quantification of internalized N-cadherin in control RNAi cells (siCtrl) and Pacsin 2 RNAi cells (siPacsin2) after normalizing the internalized N- cadherin (Pull-down) to the total amount of N-cadherin (Input) in (A). Data are means ± SD (N=3).

**Fig. 8.**
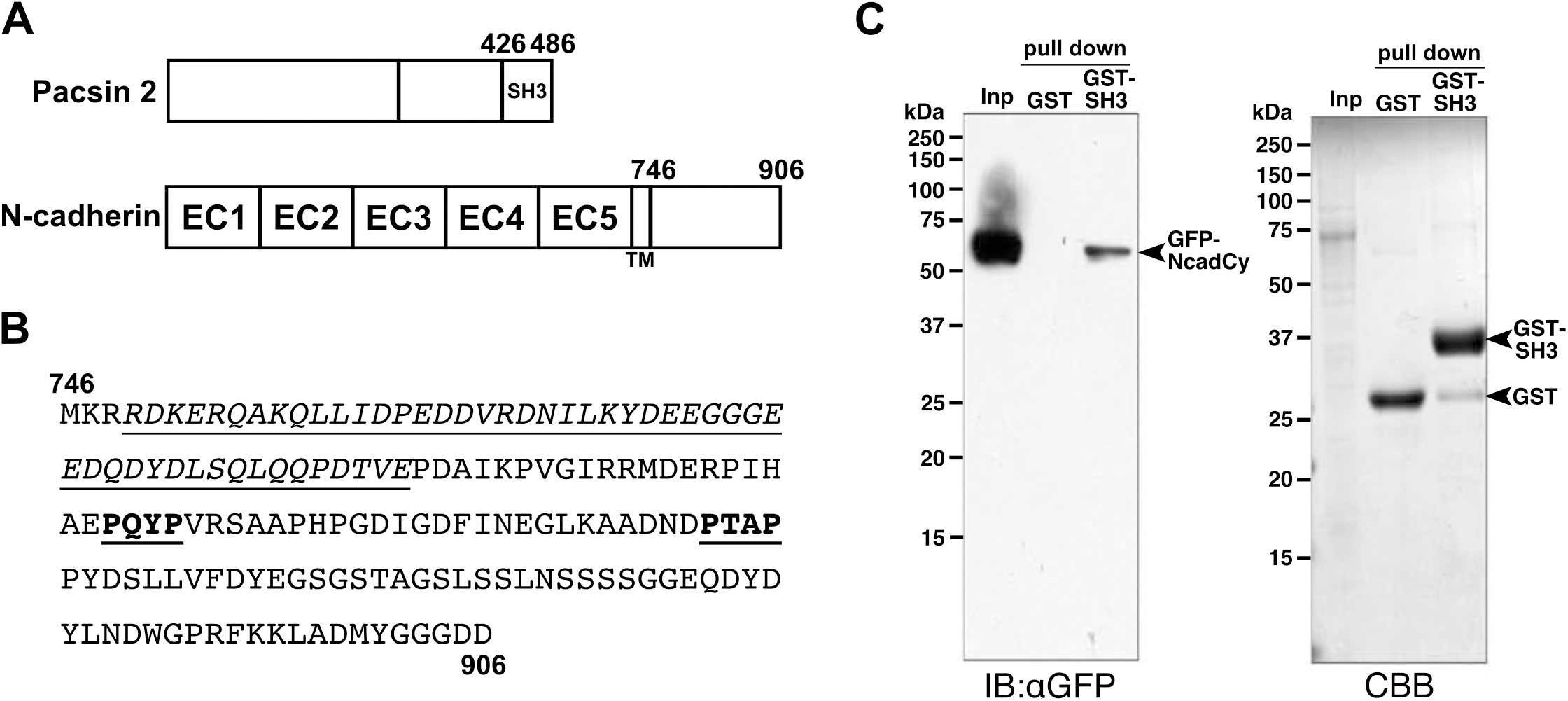
SH3 domain of pacsin 2 binds to the cytoplasmic domain of N-cadherin. (A) Schematically illustrated domain structures of human pacsin 2 and human N-cadherin. (B) Amino acid sequences of the cytoplasmic domain of N-cadherin. Two PxxP motifs (bold and underlined) and p120-catenin binding regions (italicized and underlined) are shown. (C) Interaction between pacsin 2 SH3 domain and N-cadherin cytoplasmic domain in GST pull- down assay. Immunoblot probed with anti-GFP antibody (IB: *α*GFP) and CBB-stained SDS- PAGE gel (CBB) for input (Inp) and pulled-down fraction with either GST beads (GST) or GST-pacsin 2 SH3 beads (GST-SH3) are shown. Positions for GFP-tagged N-cadherin cytoplasmic domain (GFP-NcadCy), GST (GST) and GST-tagged pacsin 2 SH3 domain (GST-SH3) are shown in arrowheads.

### Depletion of pacsin 2 and dynamin 2 enhances focal adhesion

Collective cell migration requires not only cell-cell adhesion but also integrin-based focal adhesions. A previous study showed that dynamin 2 is required for the internalization of integrins in NIH-3T3 cells (Ezratty et al., 2005). Consistently, dynamin 2 RNAi also induced an increased number of focal adhesion sites in T24 cells (19.8 focal adhesion sites per cell) compared to those in control RNAi cells (3.7 focal adhesion sites per cell) (Fig. S7, A and B). Similar to dynamin 2 RNAi cells, immunofluorescence microscopy showed that pacsin 2 RNAi cells also exhibited more than three times paxillin positive focal adhesions (18.2 - 29 focal adhesion sites per cell) compared to those in control RNAi cells (5.9 focal adhesion sites per cell) (Fig. 9, A and B). These results suggest that pacsin 2 and dynamin 2 are involved in the formation of focal adhesions as well as cell-cell contacts that are essential for collective cell migration.

**Fig. 9.**
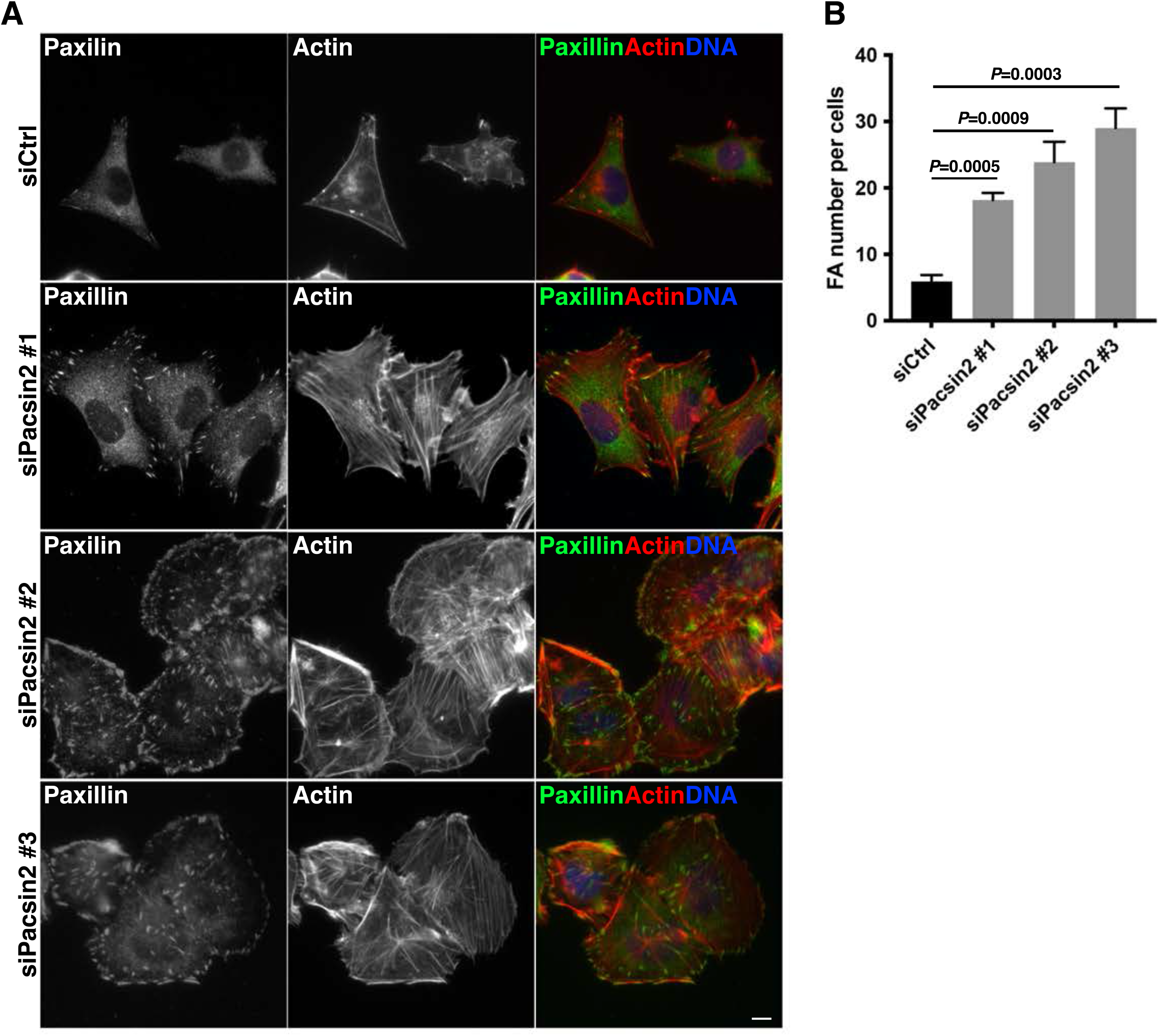
Depletion of pacsin 2 induces an elevated number of focal adhesions in T24 cells. (A) Immunofluorescence micrographs of control RNAi cells (siCtrl) and pacsin 2 RNAi cells (siPacsin 2 #1, #2 and #3) stained for a focal adhesion marker paxillin (green), F-actin (red) and their merged images with DNA (blue). (B) Quantitation of focal adhesions in control RNAi cells (siCtrl) and pacsin 2 RNAi cells (siPacsin 2 #1, #2 and #3). Data are means ± SD (n≥100 cells, N=3).

## DISCUSSION

In this study, we identified pacsin 2 as a novel regulator for collective cell migration of bladder cancer cell T24. Pacsin 2-depleted T24 cells exhibited directional migration (Fig. 2 and 3) with an increased number of cell-cell contacts enriched with N-cadherin (Fig. 4 and 5). In many epithelial cancers, metastasis is facilitated by transitioning cancer cells from epithelial to mesenchymal phenotype which is called epithelial-to-mesenchymal transition (EMT) (Nieto et al., 2016). In EMT, expression profiles of cadherin isoforms are typically switched from E- to N-cadherin so-called “cadherin switching”, which is associated with increased migratory and invasive behaviour of cancer cells (Wheelock et al., 2008). Previous studies showed that N-cadherin promotes cell aggregation and collective invasion into collagen matrices and penetration into mesothelium-like layers in lung cancer (Kuriyama et al., 2016) and ovarian cancers (Klymenko et al., 2017). Likewise, in transformed MDCK (Mardin-Darby canine kidney) cells, N-cadherin mediated cell-cell adhesion enhanced directional collective cell migration into the 3D matrix (Shih and Yamada, 2012).

Consistently, E- and N-cadherin were dominant cadherin isoforms in less aggressive (RT4) and malignant (T24) bladder cancer cells, respectively (Fig. S3) (Elie-Caille et al., 2020). Cadherin switching is controlled by either transcriptional or post-transcriptional mechanisms. In the transcriptional control, E-cadherin repression (Thiery and Sleeman, 2006) and/or N-cadherin upregulation (Maeda et al., 2005) are proposed, but the detailed mechanisms of this transcriptional regulation remain to be established. Several studies showed that E-cadherin expression level is also controlled post-transcriptionally, in which p120-catenin binds to the cytoplasmic regions of classical cadherins to prevent their degradation (Davis et al., 2003; Ireton et al., 2002; Xiao et al., 2003a). In this study, we showed that pacsin 2 RNAi or dynamin 2 RNAi altered the cell surface level of N-cadherin without affecting its total expression level (Fig. S6), suggesting that pacsin 2, as well as dynamin 2, regulates N-cadherin internalization in T24 cells to affect their migrating behaviours.

In T24 cells, pacsin 2 and dynamin 2 colocalized with N-cadherin at the cell periphery (Fig. 1 and Fig. S4). Furthermore, pacsin 2 RNAi induced attenuation of N-cadherin internalization in T24 cells (Fig. 7), suggesting that pacsin 2 is required for N-cadherin endocytosis. Previous studies showed that endocytosis of cadherins from the cell surface occurs via either clathrin-dependent or independent endocytic pathways (Cadwell et al., 2016). In MDCK cells, E-cadherin is constitutively retrieved by clathrin-dependent endocytosis (Le et al., 1999). VE-cadherin in endothelial cells is also endocytosed in a clathrin-dependent manner resulting in degradation in lysosomes (Xiao et al., 2003b). Similarly, N-cadherin is endocytosed in the clathrin-dependent pathway to facilitate neurite outgrowth (Chen and Tai, 2017). Furthermore, clathrin-dependent endocytosis is required for FGF-mediated internalization of E-cadherin (Bryant et al., 2005) and VEGF-mediated internalization of VE-cadherin (Gavard and Gutkind, 2006). A recent study using human umbilical endothelial cells showed that pacsin 2 inhibits VE-cadherin internalization from trailing ends of focal adherens junctions without affecting the total surface levels of VE-cadherin (Dorland et al., 2016). However, our study clearly showed that pacsin 2 depletion inhibits N-cadherin internalization from the cell surface (Fig. 7). Since the cytoplasmic domains of VE- and N-cadherins are divergent in their amino acid sequences, pacsin 2 may associate with multiple cadherins in various ways to control their functions required for specific cell types.

Clathrin-independent endocytic pathways are also involved in the internalization of cadherins, though it is poorly understood compared to clathrin-dependent endocytosis. Caveolin-dependent endocytosis of E-cadherin is required for disruption of cell-cell adhesion induced by EGF signalling, which is relevant to EMT of cancer cells (Lu et al., 2003). Another study showed that Rac1-modulated macropinocytosis is also required for the EGF-induced internalization of E-cadherin in breast carcinoma (Bryant et al., 2007). Clathrin- and caveolae-independent endocytosis is required for N-cadherin internalization to regulate early neuronal maturation *in vivo* (Shikanai et al., 2018). Pacsins and dynamins regulate clathrin-dependent endocytosis and caveolae-dependent endocytosis (Dessy et al., 2000; Henley et al., 1998; Qualmann and Kelly, 2000; Senju et al., 2011). Consistently, dynamin 2 RNAi cells phenocopied pacsin 2 RNAi cells in the formation of cell-cell contacts enriched with N-cadherin (Fig. S2 and S5), suggesting that pacsin 2 and dynamin 2 may cooperatively regulate the internalization of N-cadherin in clathrin-dependent and/or -independent pathways. Clathrin-mediated endocytosis of cadherins is regulated by p120-catenin, an armadillo family protein that binds to the cytoplasmic domain of classical cadherin (Reynolds, 2007). In p120- null colon carcinoma cells SW48 (Ireton et al., 2002) or p120-depleted microvascular endothelial cells MEC (Xiao et al., 2003a), cadherins are degraded through an endo- lysosomal pathway, revealing that p120 plays essential roles in the regulation of cadherin endocytosis. Interestingly, pacsin 2 SH3 domain is bound to cytoplasmic domains of N-cadherin where two PxxP motifs are located in distinct regions from p120-binding sites (Fig. 8). Thus, pacsin 2 may regulate N-cadherin endocytosis cooperatively with p120-catenin and/or in a novel mechanism independent from p120-catenin mediated endocytosis.

In this study, we also showed that depletion of either pacsin 2 or dynamin 2 induced elevated numbers of focal adhesions (FAs) in T24 cells (Fig. 9 and Fig. S7), suggesting their roles in regulating FA turnover. Indeed, previous studies demonstrated that dynamin 2 is required for FA turnover. In NIH-3T3 cells, focal adhesion disassembly induced by microtubule regrowth after nocodazole washout depends on the recruitment of dynamin 2 to FA (Ezratty et al., 2005). Another study showed that interaction between dynamin 2 and focal adhesion kinase (FAK) regulates focal adhesion dynamics in response to active Src (Wang et al., 2011). An additional study showed that the clathrin-dependent pathway is mainly involved in the dynamin 2-dependent endocytosis of FA components that lead to FA disassembly (Chao and Kunz, 2009). In contrast to dynamin 2, pacsin 2 function in FA turnover is largely unknown. Since pacsin 2 and dynamin 2 are cooperatively involved in clathrin-dependent and independent endocytosis, future studies will reveal their precise function in FA turnover.

In the collective cell migration, cell-cell and cell-extracellular matrix adhesions need to be finely balanced (Hamidi and Ivaska, 2018). Cell-cell adhesion molecules (e.g. cadherins) and focal adhesion molecules (e.g. integrins) share common signalling molecules and they are physically linked intracellularly via actin cytoskeleton (Mui et al., 2016). The convergence of crosstalk between these cell adhesion molecules is thought to be Rho family GTPases (Combedazou et al., 2020). N-cadherin in non-tumour cells enhances collective cell migration via polarization of Rho family GTPases essential for cytoskeletal regulation (Mrozik et al., 2018). N-cadherin also facilitates collective cell migration by polarizing focal adhesions in the leading cells by elevating cdc42 and Rac1 activity towards the free leading edge, resulting in enhanced cell migration (Ouyang et al., 2013; Sabatini et al., 2008; Theveneau et al., 2010). Simultaneously, Rho A is also activated at the lateral and rear sides of the leading cells inducing enhanced stress fibre formation and actomyosin contractility to establish robust cell-cell contacts (Carmona-Fontaine et al., 2008). The enhanced cell-cell contacts and focal adhesions observed in either pacsin 2 (Fig. 5 and Fig. 9) or dynamin 2 RNAi cells (Fig. S5 and Fig. S7) suggest their roles in coordinating distinct cell adhesion machinery required for coordinated movement of collectively migrating cells.

BAR domain proteins have been implicated in cancer metastasis by controlling cell motility, migration and invasion. CIP4 (Cdc42-interacting protein 4), an F-BAR domain protein, suppresses Src-induced invadopodia formation and invasion of breast cancer cells by promoting MT1-MMP endocytosis (Hu et al., 2011). I-BAR domain protein, MIM (Missing in Metastasis), suppresses metastasis by regulating cytoskeletal dynamics and lamellipodia formation, consequently affecting the invasion and metastatic behaviour of cancer cells (Woodings et al., 2003). Furthermore, an N-BAR domain protein endophilin regulates endocytosis of EGFR by controlling F-actin cytoskeleton (Vehlow et al., 2013). Interestingly, expression of brain-specific isoform pacsin 1 is negatively correlated with the malignancy of gliomas, indicating that pacsin 1 would play an essential role in the development of gliomas and might be a potential new biomarker and targeted therapy site for gliomas (Zimu et al., 2021). Based on the TCGA PanCancer Atlas studies in cBioPortal, either deep deletion or mutations in pacsin 2 gene were identified in patients of different types of malignant cancers including ovarian cancer, breast cancer and bladder cancer (Cerami et al., 2012; Gao et al., 2013). Thus, future studies about the correlation between the expression level of pacsin 2 and the malignancy of various cancers may bring pacsin 2 as a potential therapeutic target in cancer metastasis.

## MATERIALS AND METHODS

### Molecular biology

Expression constructs were prepared using Gateway Cloning (Thermo Fisher Scientific) as described previously (Fujise et al., 2021). To prepare Entry clones for either N-cadherin cytoplasmic domain or Pacsin 2 SH3 domain, PCR fragments amplified from clones of human N-cadherin or Pacsin 2 using primers described in the supplementary information was used for B–P recombination with either pDONR201 (Pacsin 2 SH3) or pENTR/D-TOPO (N-cadherin cytoplasmic domain). These Entry clones were subcloned into pCI based Destination vectors for expressing GFP-tagged proteins in mammalian cells (N-cadherin cytoplasmic domain) or GST-tagged proteins in bacterial cells (Pacsin 2 SH3) by L-R recombination.

### Cell culture, DNA transfection, and RNAi

T24 cells (ATCC HTB-4) were cultured in RPMI-1640 medium (189-02025, FUJIFILM Wako Chemicals) supplemented with 10% fetal bovine serum (FBS) (26140-079, Gibco) and penicillin-streptomycin (PS) (100 unit/ml) (15140122, Thermo Fisher Scientific) at 37°C in humidified air with 5% CO_2_. HEK293T cells (ATCC CRL-3216) were grown in D-MEM (High Glucose) with L-Glutamine, Phenol Red, and Sodium Pyruvate (043-30085, FUJIFILM Wako chemicals) supplemented with 10% FBS (26140-079, Gibco) and PS (100 unit/ml) (15140122, Thermo Fisher Scientific) at 37°C in 5% CO_2_.

To transfect HEK293T, 70% of confluent cells in VIOLAMO VTC-P6 6-well plates (2- 8588-01, AS ONE) were transfected with 1.5 μg expression plasmids using Lipofectamine LTX with Plus Reagent (15338-100, Thermo Fisher Scientific) following manufacturer’s instructions. The cells were collected 48 h after the transfection and used for GST pulldown assay.

For RNAi of T24 cells, 70% confluent cells in 24-well plates were transfected with 10 pmol of either siGENOME SMART pool siRNA for human Dnm2 (M-04007-03, Dharmacon) or siGENOME nontargeting siRNA Pools #1 (D-001206-13-05, Dharmacon), Mission siRNA for human Pacsin 2 siRNA (SASI_Hs01_0021-5538, SASI_Hs01_0021-5539 and SASI_Hs01_0021-5540, Merck) or MISSION siRNA Universal Negative Control #1 (SIC-001, Merck) using Lipofectamine RNAiMAX Transfection Reagent (13778150, Thermo Fisher Scientific) following manufacturer’s instructions.

### Antibodies and reagents

Primary antibodies used in this study were rabbit polyclonal anti-dynamin 2 (ab 65556 Abcam), rabbit polyclonal anti-PACSIN1 (M-46) (SC-30127, Santa Cruz), mouse monoclonal anti- PACSIN 2 (SAB1402538-100UG, SIGMA), rabbit polyclonal anti-PACSIN3 (AB37612, Abcam), mouse monoclonal anti E-cadherin (610181, BD Transduction laboratory), mouse monoclonal anti N-cadherin(610920, BD Transduction laboratory), mouse monoclonal anti P-cadherin (12H6, Cell Signaling Technology), mouse monoclonal anti VE-cadherin (610251, BD Transduction laboratory), mouse monoclonal anti-Paxillin (5H11) (AH00492, Thermo Fisher Scientific), mouse monoclonal anti-alpha tubulin (T5168, Merck), rabbit polyclonal anti-cortactin (A302-608A-M, Thermo Fisher Scientific) and GFP (D5.1) XP rabbit mAb (2956, CST). Secondary antibodies and Alexa-conjugated phalloidin for immunofluorescence microscopy were purchased from Thermo Fisher Scientific: Alexa Fluor 488 donkey anti-goat IgG (H+L) (A-11055), Alexa Fluor 488 donkey anti-rabbit IgG (H+L) (A-21206), Alexa Fluor 555 donkey anti-mouse IgG (H+L) (A-31570), Alexa Fluor 555 donkey anti-rabbit IgG (H+L) (A-31572) and Alexa Fluor 555 Phalloidin (A-34055). Secondary antibodies for immunoblot analyses were also purchased from Thermo Fisher Scientific: goat anti-rabbit IgG (H+L) secondary antibody, HRP (31460), rabbit anti-mouse IgG (H+L) secondary antibody, HRP (31450).

PFA used for cell fixation was prepared from 16% paraformaldehyde (15710, Electron Microscopy Sciences). Glutaraldehyde used for fixing gelatin-coating coverslips was prepared from Glutaraldehyde 25% EM (G004, TAAB).

### Immunofluorescence microscopy

Immunofluorescence microscopy of T24 cells on the gelatin-coated coverslips was performed as described previously (Li et al., 2021) using primary antibodies (1:100 for E-cadherin and N-cadherin antibodies, 1:200 dilution for Dynamin2, Cortactin, Paxillin and Pacsin 2 antibodies) and secondary antibodies (1:1000 dilution) and Alexa Fluor 555 Phalloidin (1:1000 dilution). Immunostained samples for T24 cells were visualized using BX51 fluorescence microscope (OLYMPUS) with 40 × NA 0.75 objective lens and images were acquired with Discovery MH15 CMOS camera (Tucsen) and ISCapture image acquisition software (Tucsen). All images were analyzed using Fiji (Schindelin et al., 2012) and processed with Adobe Photoshop 2022 (Adobe).

### Immunoblot analysis

Immunoblot analyses were performed as described previously (Fujise et al., 2021) using primary antibodies (1:1000 dilution) and secondary antibodies (1:10000 dilution).

### Wound healing assay

Confluent T24 cells transfected with control and pacsin 2 RNAi in 6-well plates were treated with 10 ng/ml of recombinant human EGF (236-EG, R&D systems) in RPMI-1640 medium and incubated at 37℃, 5% CO_2_ for 2 hours. After 2 hours, the cell monolayer was scratched with a sterile 200 μl pipette tip. Recovery processes were captured after 0, 6 and 12h using a NEX-5N camera (Sony) attached to Nikon Eclipse Ts2 microscope using a 2× objective lens.

### Live-cell imaging of wound healing assay

For live cell imaging of the wound healing assay, T24 cells (0.6×10^6^ cells) transfected with either control siRNA or pacsin 2 siRNA were seeded in a 35 mm Glass Base Dish (3910-035, IWAKI) and incubated at 37℃, 5% CO_2_ until the cell monolayer became confluent. Cells were then treated with 10 ng/ml of recombinant human EGF (236-EG, R&D systems) in RPMI-1640 medium and incubated at 37℃, 5% CO_2_ for 2 hours. After the cell monolayer was scratched with a sterile 20 µl pipette tip, cells were maintained in 5% CO_2_ at 37℃ with a thermo-control system (MI-IBC, OLYMPUS) and images were acquired on IX71 microscope (OLYMPUS) fitted with X-Light spinning disc confocal unit (CrestOptics) and iXon EMCCD camera (DU- 888E-C00-#BV, ANDOR) using MetaMorph (Molecular Devices). Images were captured with a 20× objective lens (LCPlanFI 20x/0.40) at 1 minute intervals for 6 hours.

For cell tracking, ten cells at the front of either side of the wound edge were tracked by Manual Tracking plugins of Fiji (Schindelin et al., 2012). Cell velocity and directionality were determined by analyzing the trajectories of each cell by Chemotaxis Tool (ibidi). Cell trace was analyzed by OriginPro 2021 (Origin Lab).

### GST pull-down assays

Recombinant protein of human pacsin 2 SH3 domain (aa 426–486) was expressed and purified as GST fusion using Glutathione Sepharose 4B (17075601, Cytiva) according to the manufacturer’s instructions to be used as bait for the GST pull down assay. GFP-tagged cytoplasmic domain of N-cadherin (aa 746-906) was expressed in HEK293T cells and the cell extract was prepared by sonication using TAITEC VP-5S sonicator (output: 5; 5 s × 3 times) in extraction buffer (20 mM Tris-HCl, pH 7.5, 150 mM NaCl, 10% glycerol, 5 mM EDTA, 2% Triton X-100, 1 mM PMSF and protease inhibitor (11697498001, Roche)) followed by centrifugation at 20,600×g for 10 min at 4℃. For the pulldown assay, 170 µl of the cell extract was added to 10 µl (bed volume) of GST-Pacsin 2 SH3 or GST beads in the extraction buffer and mixed for 1 hr at 4°C with gentle agitation. After washing three times with the extraction buffer, the bound proteins were analyzed by immunoblot analyses.

### Surface biotinylation and endocytosis assay

Surface biotinylation and endocytosis assay was performed as described previously (Morimoto et al., 2005) with minor modifications. T24 cells (2.5x 10^5^ cells) transfected with control siRNA or pacsin 2 siRNA for 48 hours were seeded in a 100 mm TC-treated culture dish (430167, Corning) in RPMI-1640 medium. After 24 hours, cell surface proteins were biotinylated with 0.5 mg/ml Ez-link sulfo-NHS-SS-Biotin (21331, Thermo Fisher Scientific) in PBS containing 1 mM CaCl_2_ and 0.5 mM MgCl_2_ (PBS/CM) at 4°C for 30 min, and quenched with 50 mM NH_4_Cl in PBS/CM at 4 °C for 15 min. The cells were then allowed endocytosis at 37°C for the indicated periods until the endocytosis was stopped by rapid cooling of the cells on ice at 4°C. The remaining biotin on the cell surface was stripped with 50 mM of sodium 2-mercaptoethanesulphonate (MESNa) (M1511, Sigma-Aldrich) in PBS/CM at 4°C for 30 min followed by quenching with 5 mg/ml iodoacetamide (093-02152, Fujifilm) in PBS/CM at 4°C for 15 min. Cells were then lysed with RIPA buffer (20 mM Tris-HCl pH-7.6, 150 mM NaCl, 0.1% SDS, 1% Triton X-100, 1% deoxycholate, 5 mM EDTA with protease inhibitors (11697498001, Roche)) by sonication using TAITEC VP-5S sonicator (output: 5; 5 s × 2 times) and cell lysate was recovered in the supernatant after centrifugation at 20,000×g at 4°C for 15 min. The cell lysates containing an equal amount of total proteins (≃2 mg in 1ml RIPA buffer) were incubated with 15 µl (bed volume) of Pierce NeutrAvidin agarose (29200, Thermo Fisher Scientific) for 2 hours at 4°C to capture internalized biotinylated protein. After washing three times in RIPA buffer, biotinylated proteins were eluted from NeutrAvidin agarose beads by sample buffer, separated by SDS-PAGE and analyzed by immunoblotting using an antibody against N-cadherin. The relative amount of internalized N-cadherin (Pull-down) was obtained by normalizing to the total amount of N-cadherins expressed in the cells.

### Electron microscopy

T24 cells were fixed in 2% glutaraldehyde and 2% paraformaldehyde at 4°C overnight and then postfixed in 2% osmium tetraoxide for 1.5 h at 4°C followed by dehydration with ethanol and embedding in Spurr resin (Polysciences Inc., Warrington, PA). Ultrathin sections of the samples were prepared using Ultramicrotome (Leica EM UC7) and they were stained with uranyl acetate and lead and observed by electron microscope H-7650 (Hitachi, Tokyo, Japan) at Central Research Laboratory (Okayama University Medical School).

### Statistical analysis

All the experiments were repeated three times independently. Data were analyzed for statistical significance using GraphPad Prism6 (GraphPad Software).

## Acknowledgements

This work was supported by JSPS KAKENHI, Grant numbers 18K07198, 19KK0180, grants from Wesco Scientific Promotion Foundation for T.T., 19H03225 for K.T., and 19H01064 for M.W. The authors also thank the technical assistance of Ms Masumi Furutani and Ms Rumi Tsukano (Okayama University Medical School) for their technical assistance in electron microscopy.

## Notes

### Competing Interest Statement

The authors have declared no competing interest.

