## Supplementary Files for "Pacsin 2-dependent N-cadherin internalization regulates the migration behaviour of malignant cancer cells"

#### *SUPPLEMENTARY INFORMATION*

*Primers*

**Pacsin 2 SH3:**

5'-GGGG ACA AGT TTG TAC AAA AAA GCA GGC TGC gggacggaagtgcga-3'

5'-GGGG ACC ACT TTG TAC AAG AAA GCT GGG T tcactggatcgcctcc-3'

**N-cadherin cytoplasmic domain**

5'-CACCATGAAACGCCGGGATAAAG-3'

5'-TCAGTCATCACCTCCACC-3'

#### *SUPPLEMENTARY FIGURES*

***Fig. S1. Expression profiles and subcellular localization of pacsin isoforms in T24 cells***. (A) Immunoblot analysis of endogenous pacsin 1, pacsin 2 and pacsin 3 in T24 cells. (B) Localization of endogenous pacsin 1, pacsin 2 and pacsin 3 (green), F-actin (red), and their merged images with DNA (blue). Scale bar is 10 μm.

***Fig. S2. Depletion of dynamin 2 induces cell-cell contacts in T24 cells.*** (A) Immunofluorescence micrographs of F-actin (red) and its merged images with DNA (blue) in control RNAi (siCtrl) or dynamin 2 RNAi (siDNM2) cells. The scale bar is 10 μm. (B) Quantitation of relative number of cells with cell-cell contacts for control RNAi cells (siCtrl) or dynamin 2 RNAi cells (siDNM2). Data are means ± SD (n≥130 cells, N=3)

***Fig. S3. Expression profiles of cadherins in T24 cells.*** Immunoblot analysis of endogenous E-,

N-, P- and VE-cadherin in RT4 or T24 cells.

***Fig. S4. N-cadheirn colocalizes with pacsin 2 and dynamin 2 at the cell periphery in T24 cells.*** Immunofluorescence micrographs of endogenous pacsin 2 (green) with endogenous N-cadherin (red) and their merged images (upper panel) or endogenous N-cadherin (green), endogenous dynamin 2 (red) and their merged images (lower panel). Scale bar is 10 µm.

***Fig. S5. Depletion of dynamin 2 induces N-cadherin-rich bridges between contacting cells.*** (A) Immunofluorescence micrographs of control RNAi cells (siCtrl) or dynamin 2 RNAi cells (siDNM2) stained for endogenous N-cadherin (green), F-actin (red) and their merged images with DNA (blue). Scale bar is 10 µm.

***Fig. S6. Expression level of N-cadherin in T24 cells is not affected by depletion of either pacsin 2 or dynamin 2.*** (A) Immunoblot analyses of cell extract from either control RNAi cells (siCtrl) or pacsin 2 RNAi (siPacsin 2 #1, #2 and #3) cells using antibodies against N-cadherin (IB: N-cadherin), pacsin 2 (IB: Pacsin 2) or $\alpha$Tubulin (IB: Tubulin) as an internal control. Quantitation of N-cadherin levels relative to α-tubulin in control RNAi cells (siCtrl) or pacsin 2 RNAi cells (siPacsin 2 #1, #2 and #3) are also shown. Data are means ± SD (N=3) (B) Immunoblot analysis of cell extract from in either control RNAi cells (siCtrl) or dynamin 2 RNAi (siDNM2) cells using antibodies against N-cadherin (IB: N-cadherin), dynamin 2 (IB: dynamin 2) or $\alpha$Tubulin (IB: Tubulin) as an internal control. Quantitation of N-cadherin levels relative to α-tubulin in control RNAi (siCtrl)or dynamin 2 RNAi (siDNM2) cells are also shown. Data are means ± SD (N=3)

***Fig. S7. Depletion of dynamin 2 induces enhances focal adhesion in T24 cells.*** (A) Immunofluorescence micrographs of Paxillin (green), F-actin (red) and their merged images with DNA (blue) in control RNAi cells (siCtrl) and dynamin 2 RNAi cells (siDNM2). (B) Quantitation of focal adhesions in control RNAi cells (siCtrl) and dynamin 2 RNAi cells (siDNM2). Data are means ± SD (n≥120 cells, N=3).

***Movie S1. Control RNAi cells in the wound healing assay.*** Images were obtained every 1min for 6 hours after the start of the wound healing assay. Trajectories of the ten representating cells shown in different colours.

Movie S2. ***Pacsin 2 RNAi cells in the wound healing assay.*** Images were obtained every 1min for 6 hours after the start of the wound healing assay. Trajectories of the ten representating cells shown in different colours.

### Fig.S1 Wint et al

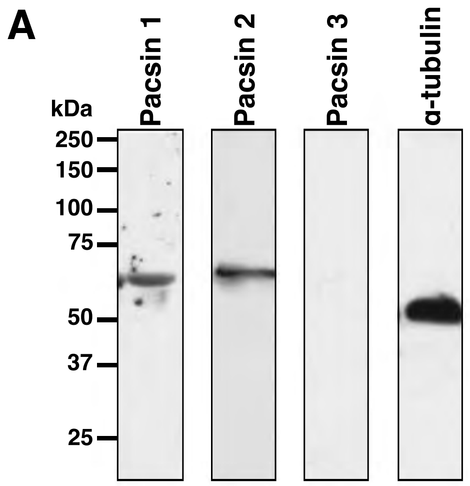


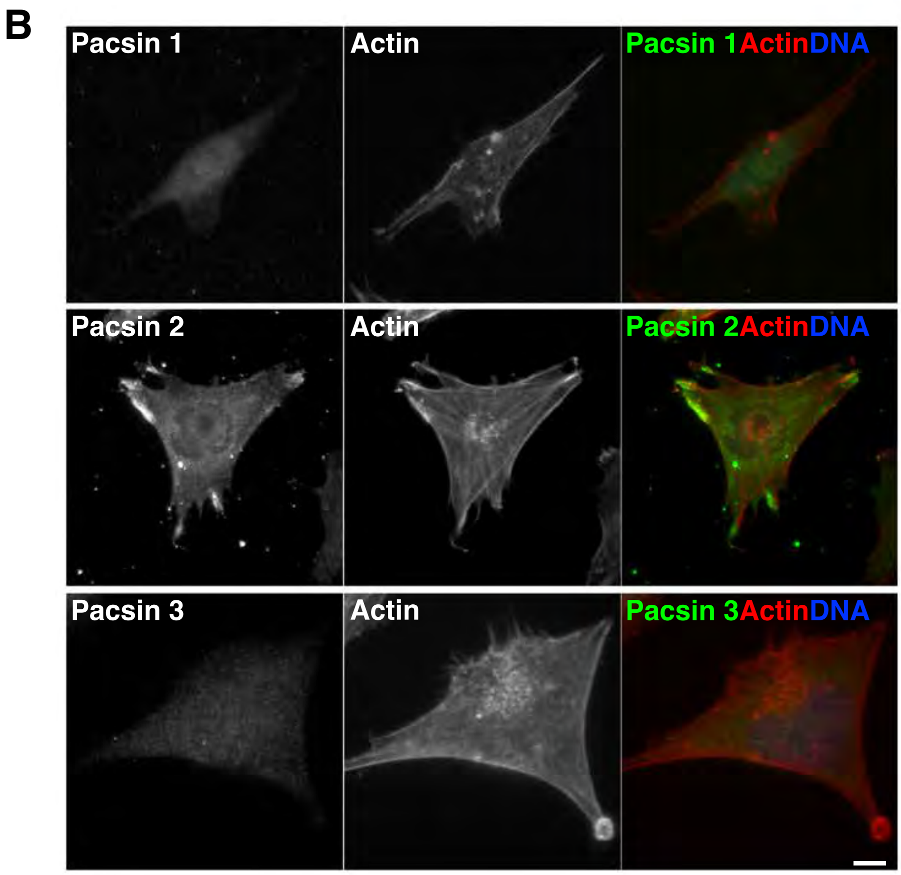

**Fig. S2 Wint et al**

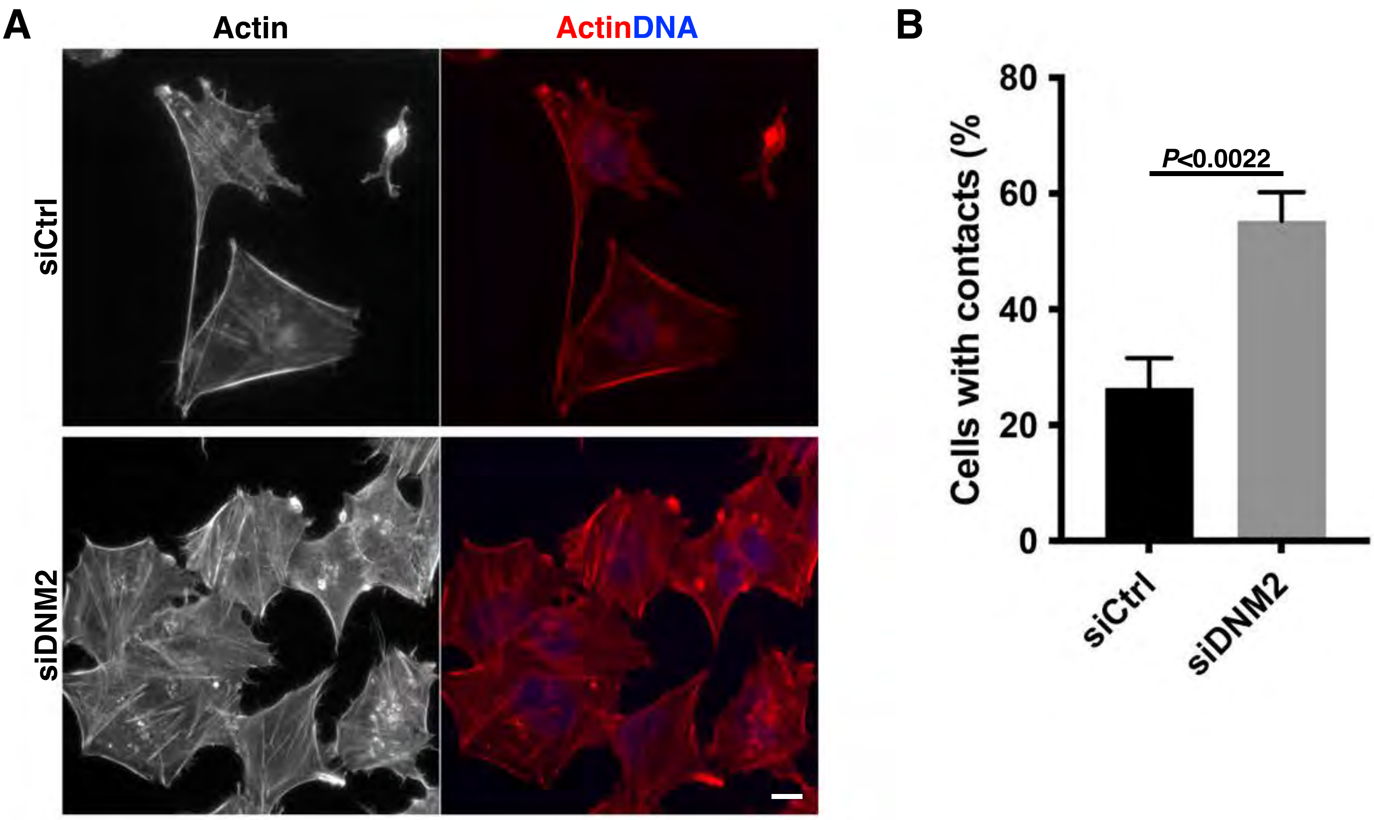

### Fig.S3 Wint et al

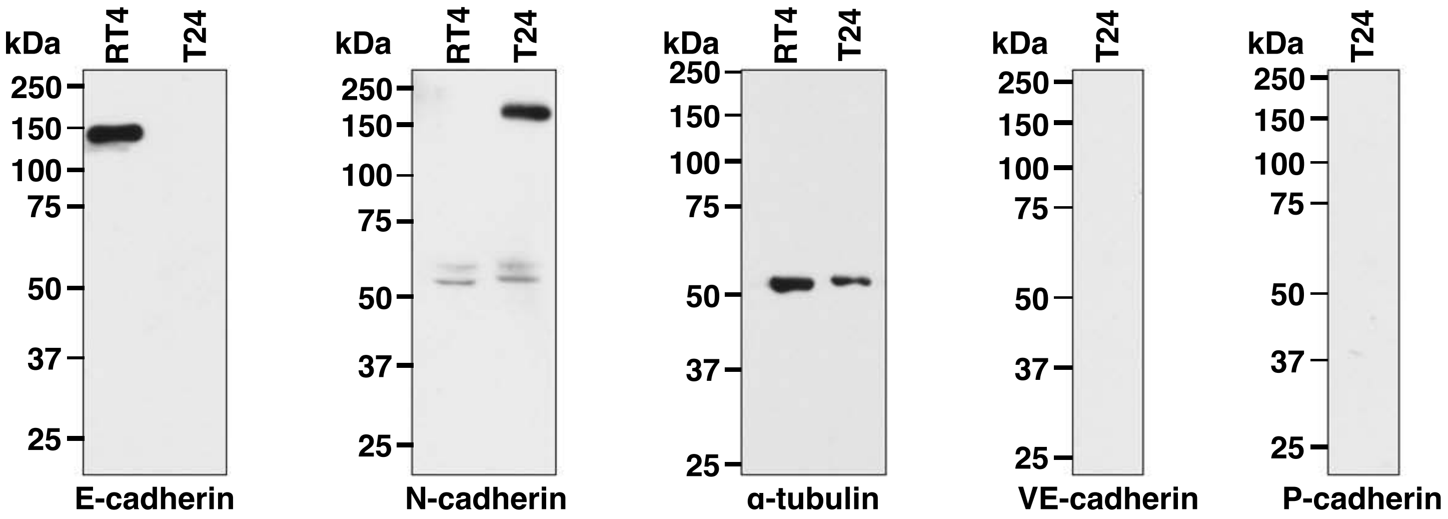

### Fig.S4 Wint et al

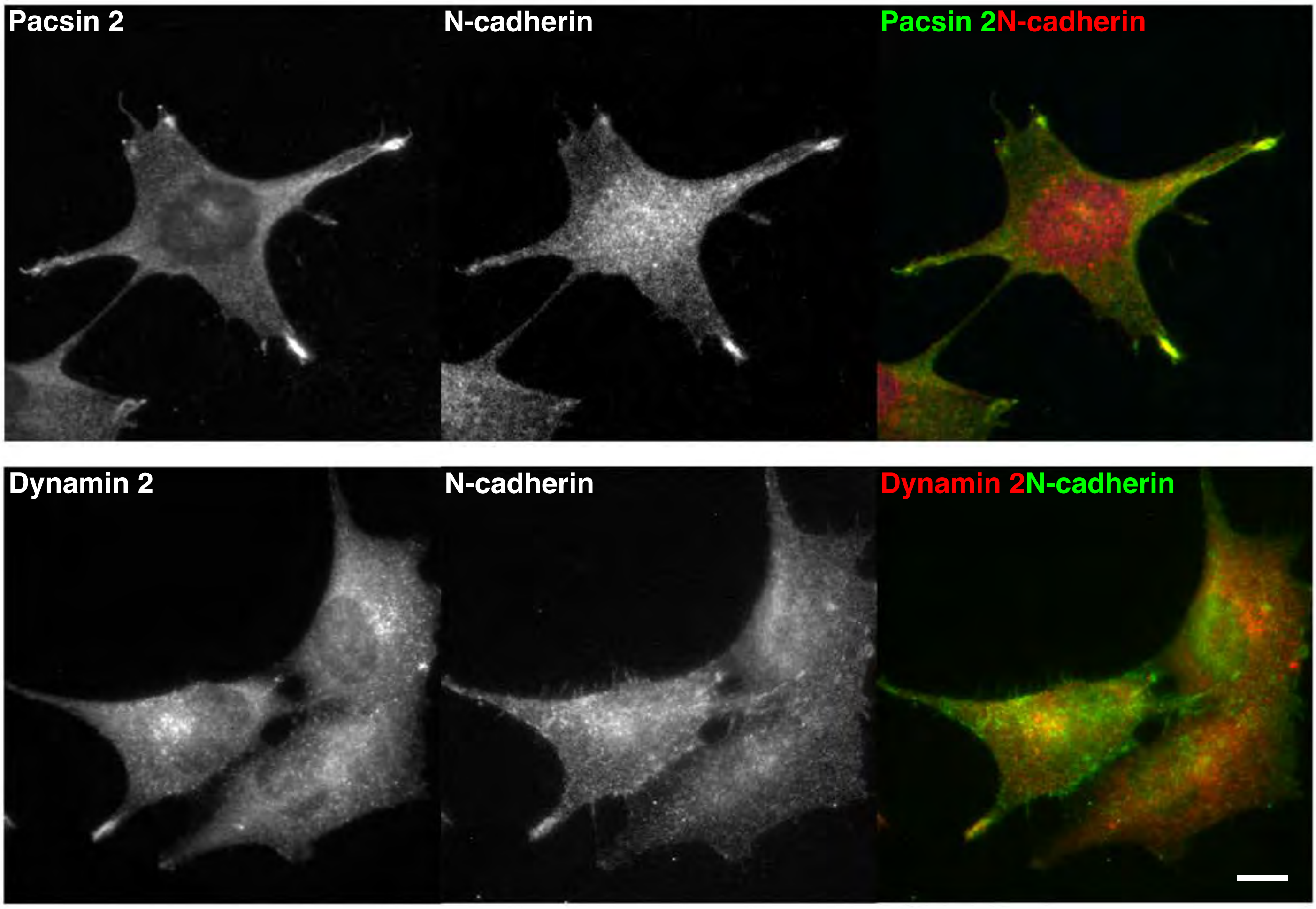

### Fig.S5 Wint et al

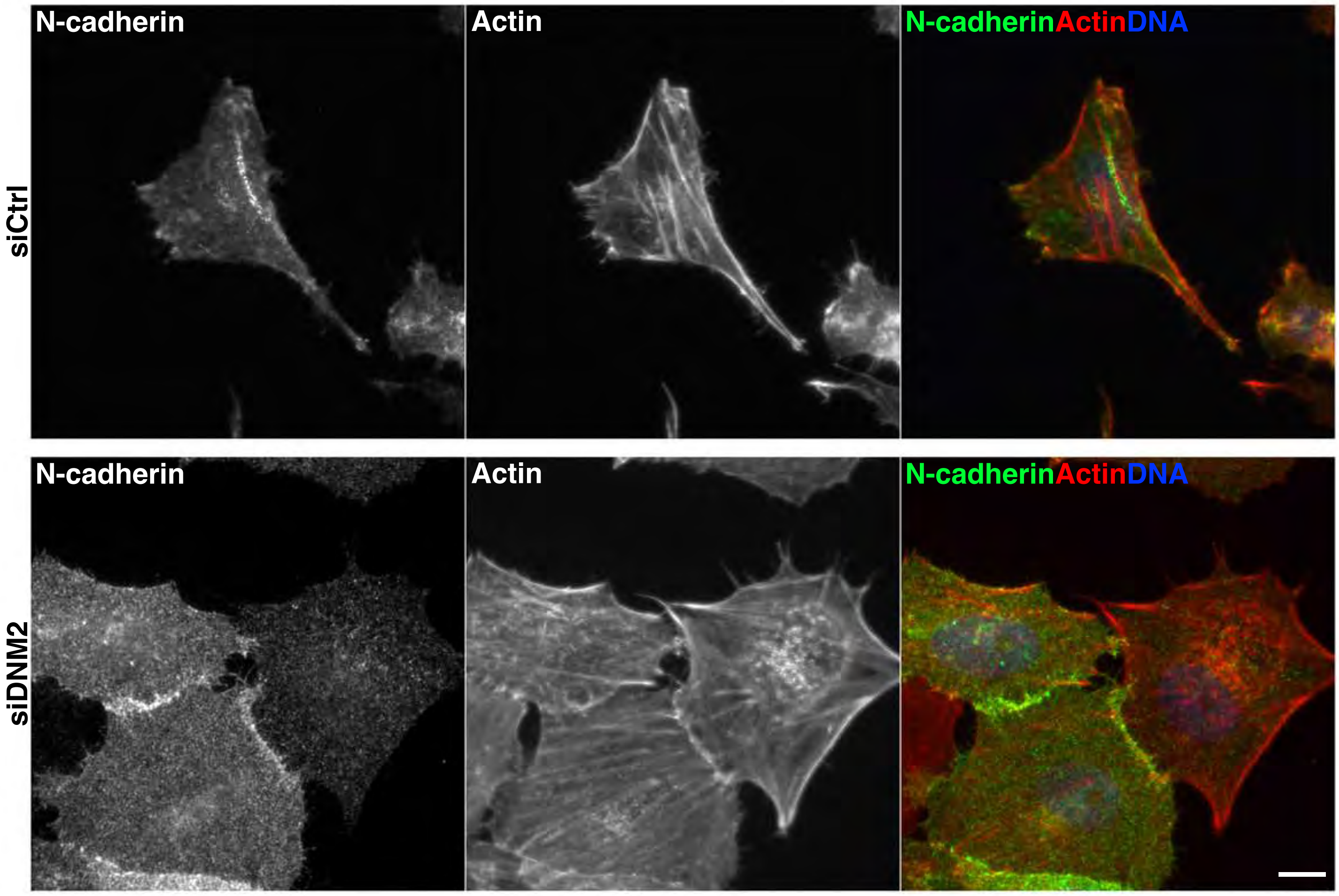

### Fig. S6 Wint et al., 2022

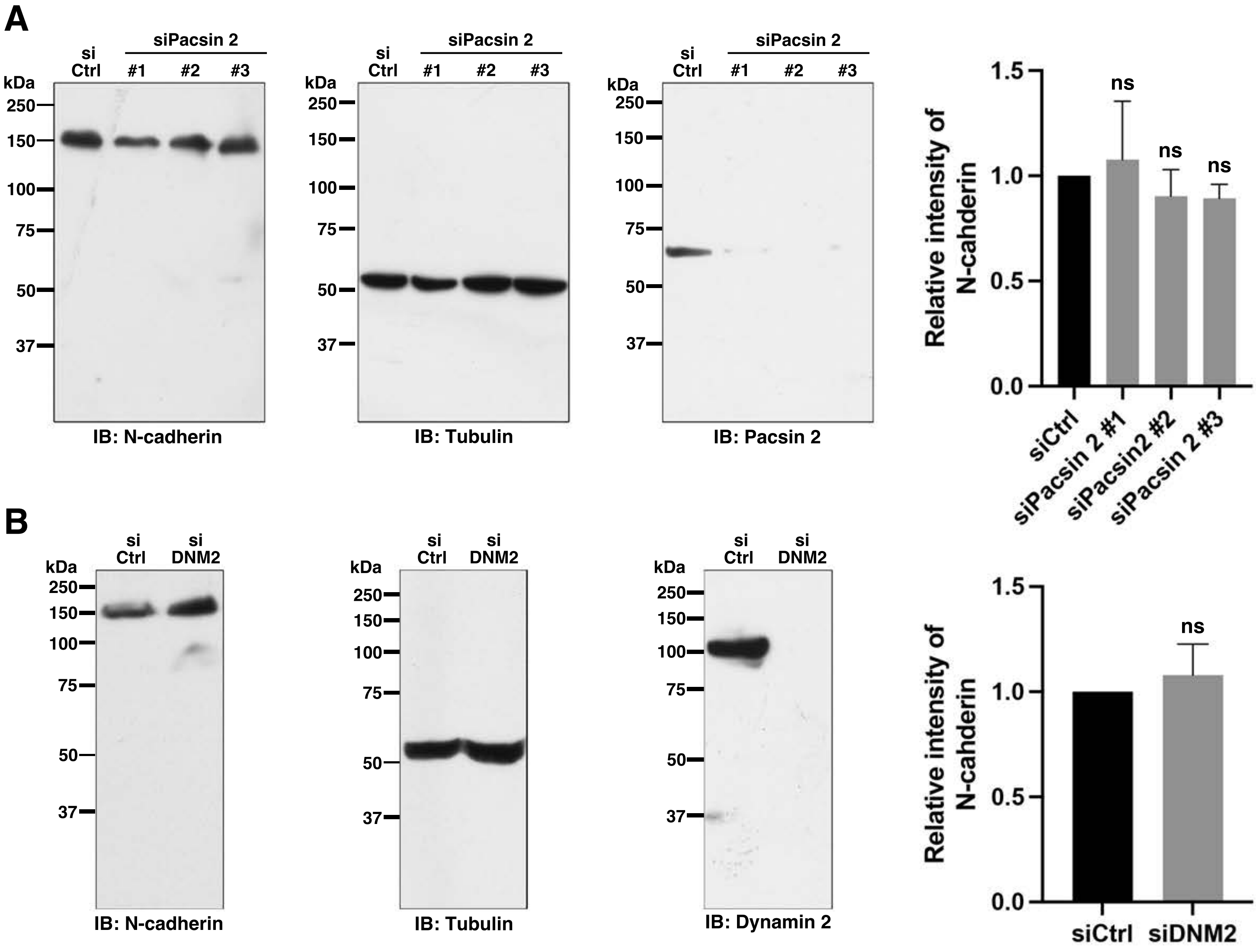

### Fig.S7 Wint et al

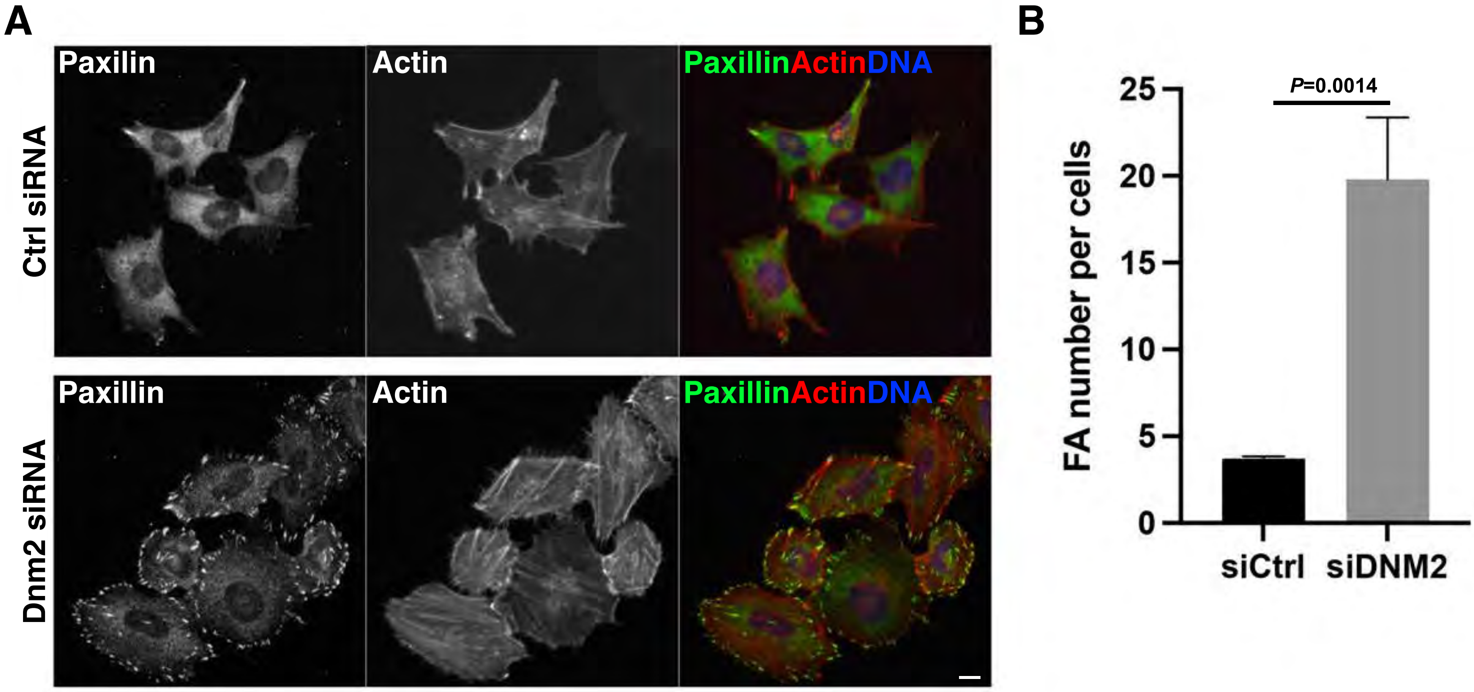
